# Characterisation of immune cell subsets of tumour infiltrating lymphocytes in brain metastases

**DOI:** 10.1101/2021.02.10.430531

**Authors:** Priyakshi Kalita-de Croft, Haarika Chittoory, Tam Nguyen, Jodi M Saunus, Woo Gyeong Kim, Amy McCart Reed, Malcolm Lim, Xavier M De Luca, Kaltin Ferguson, Colleen Niland, Roberta Mazzieri, Riccardo Dolcetti, Peter T Simpson, Sunil R Lakhani

## Abstract

The heterogeneity of tumor infiltrating lymphocytes is not well characterized in brain metastasis. To address this, we performed a targeted analysis of immune cell subsets in brain metastasis tissues to test which immunosuppressive routes are involved in brain metastasis. We performed multiplex immunofluorescence (mIF), using commercially available validated antibodies on twenty formalin-fixed paraffin embedded whole sections. We quantitated the subsets of immune cells utilizing a targeted panel of proteins including PanCK, CD8, CD4, VISTA and Iba1, and analyzed an average of 15000 cells per sample. We classified tumours as either high (>30%) or low (<30%) tumour infiltrating lymphocytes (TILs) and found that increased TILs density correlated with survival. We next sought out to phenotype these TILs using mIF. The tumours with low TILs (n=9) had significantly higher expression of the immune-checkpoint molecule VISTA in tumor cells (p<0.01) as well as in their microenvironment (p<0.001). Contrastingly, the brain metastatic tumours with high TILs (n=8) displayed higher levels of activated microglia, as measured by Iba1 expression. Low TILs-tumours displayed CD8+ T-cells that co-express VISTA (p<0.01) significantly more compared to high TILs group, where CD8+ T-cells significantly co-express Iba1 (p<0.05). Interestingly, no definite phenotypes of CD4+ subsets were observed. These results were supported by RNA analysis of a publicly available, independent cohort. In conclusion, our work contributes to a growing understanding of the immune surveillance escape routes active in brain metastasis.

**Graphical Abstract:** Graphical abstract:
Brain metastasis patients with low TILs have high VISTA expression and patients with high TILs have significantly more activated microglia

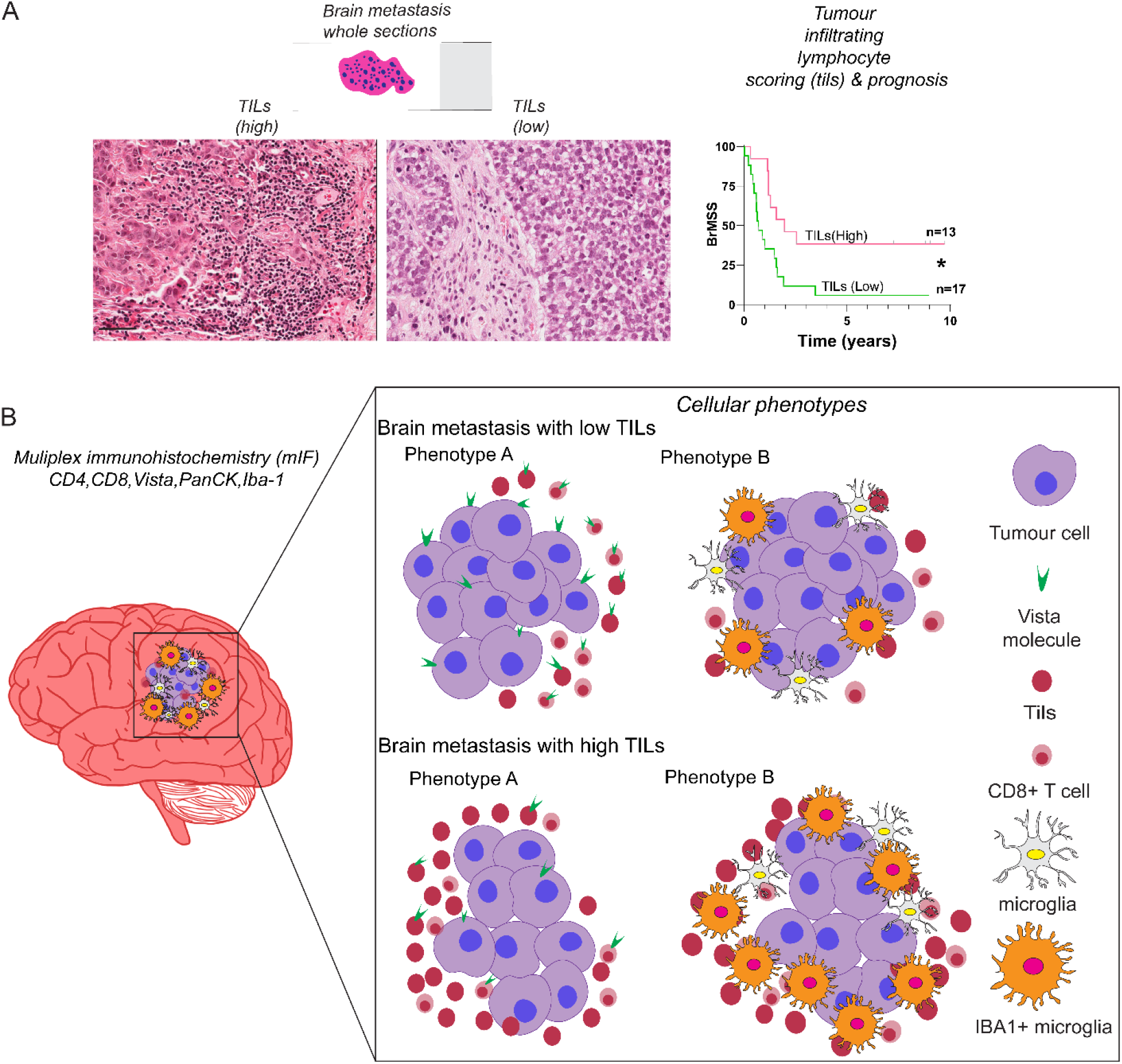

## 1. Introduction

Metastatic tumours to the brain, which originate from extracranial primaries such as melanoma, breast and lung remain a significant clinical challenge. Life expectancy of patients after being diagnosed with brain metastases (BrM) can be measured in months [1] and BrMs make up a significantly higher proportion of adult brain tumors compared to primary brain tumours. Currently the gold standard for treating BrMs is surgery, chemotherapy, and radiotherapy. The advent of immunotherapies and their proven efficacy in a subset of BrM patients [2–4] has provided new hope for further therapeutic intervention. However, treatment alternatives still remain limited due to the lack of our comprehensive understanding of the heterogeneity of the microenvironment of the brain. This brain tumor microenvironment (TME) comprises various cell types that can regulate progression of the cancer and also response to therapy [5]. Given the unique adaptations of the brain TME [6–11] as well as the presence of uncommon cell types; microglia (MG), neurons, and astrocytes; and, the blood-brain barrier, makes it compelling to further address the critical question of how TME heterogeneity affects therapeutic efficacy.

Recent work has compared the immune landscape of BrMs and gliomas, where distinctive tumor specific features were discovered [12,13]. These ground breaking studies observed more prominent infiltration of lymphocytes and leukocytes in BrMs compared to gliomas. However, targeted interrogation of the subtypes may be helpful in delineating prognostic cell subsets and specific phenotypes of these infiltrating lymphocytes. In light of this biological question, we explored the infiltrating lymphocyte phenotypes of BrMs arising from breast cancers. We undertook a targeted approach of investigating five markers of interest; Pan cytokeratin (PanCK), CD4, CD8, V-domain immunoglobulin suppressor of T-cell activation (VISTA), Iba-1 to elucidate the immune cell subtypes of BrMs that stratify according to the density of the tumor infiltrating lymphocytes (TILs). We utilized multiplex Immuno-Fluorescence (mIF) to address a number of questions in whole BrM sections. Do tumours with low density of TILs display a different phenotype compared to tumor with high density of TILs? Do the T-cell subtypes display any exhausted phenotypes? Does the density of the microglia differ in these tumours? We focused on the expression of the T-cell inhibitor molecule VISTA for our study as it is largely unexplored in BrMs from breast cancers.

VISTA was first identified by Wang and colleagues [14], where they discovered VISTA to have homology to the extracellular domain of B7 ligand programmed death ligand 1 (PD-L1). They found VISTA’s expression to be highly regulated on the myeloid antigen presenting cells and this inhibited T-cell proliferation and cytokine production. Since this discovery, knowledge on the role and function of VISTA in many cancers have been uncovered[15–19]. Furthermore, recent reports on mouse models of brain metastasis suggest VISTA blockade along with anti-PD L1 reduces BrM outgrowth and additionally, the release of cytokine Cxcl10 from tumor cells results in recruitment of VISTA expressing myeloid cells leading to T-cell suppression [20].

We interrogated the tumor-intrinsic as well as immune microenvironment specific expression of our marker panel. By exploring a targeted panel of markers within this selected cohort, we discovered that patients with reduced TILs density have increased VISTA+ immune cells as well as tumour cells. This finding enhances our understanding of the immune-suppressive phenotypes of BrM patients with high and low density of TILs. Our study provides a clinical perspective in understanding the association of VISTA expression on the tumor as well as the microenvironment compartment of BrMs using whole tissue sections. This, in turn, may help us uncover new promising, immunotherapies, as well as answer fundamental biological questions related to BrM’s immunosuppressive TME.

## 4. Materials and Methods

### Ethics

Ethical approval from the Human Research Ethics Committees of the Royal Brisbane and Women’s Hospital (RBWH; 2005000785) and The University of Queensland (HREC/2005/022) were obtained prior to the commencement of this study and de-identified samples were used for all the analyses performed.

We compiled a cohort of brain metastases arising from breast cancers undergoing resection at the Royal Brisbane and Women’s hospital, Brisbane, Australia (n=36). Based on the amount of necrosis and tissue availability, we performed multiplex immunofluorescence (mIF) on 20 whole tissue sections from 20 patients.

### TILs scoring

Immune cell infiltration was scored within the boundary of the tumor by a Pathologist (WGK). As there is no set diagnostic criteria for scoring TILs in brain metastases, for our research purposes we adapted the Guidelines for TILs assessment from the “International Immuno-Oncology Biomarker Working Group” [21]. Immune infiltrate was quantitated by the area occupied by mononuclear inflammatory cells (lymphocytes and plasma cells) over total stromal area. TILs within the borders, invasive edges and desmoplastic stroma of the metastatic tumors were included in the evaluation. The proportion of TILs within the stromal area was measured in percentage and initially criteria was set as per the guidelines. However, we restratified this criteria to > or < 30% TILs as this showed the best stratification within our cohort.

### Transient knockdown

MDA-MB-468 cells were purchased from the American Type Culture Collection (Manassas, VA, USA) and grown under standard culture conditions. Transient knockdowns were performed with mycoplasma free cells using 100 nM siRNAs comprising of three different probes (Gene Pharma, Shanghai, China) and FugeneHD (Promega, Madison, WI, USA) for 24 hours. The cells were fixed in 10% neutral buffered formalin (Sigma-Aldrich, Missouri, USA) after 24 hours and embedded in paraffin for antibody validation by immunohistochemical analysis.

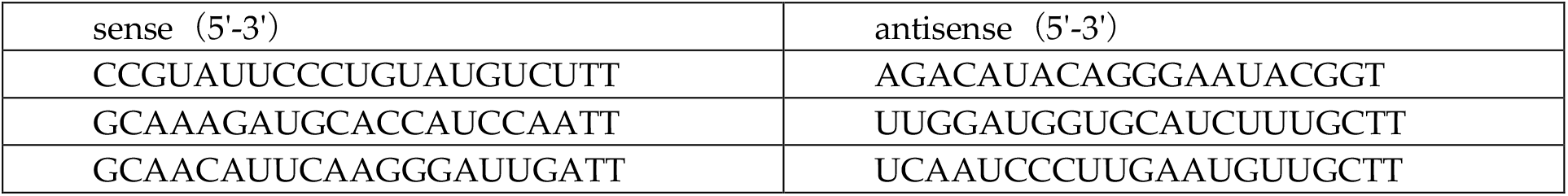

### mIF and IHC

Four-micron thick sections were used to perform mIF for detecting CD4, CD8, Vista, PanCK and Iba-1 using Tyramide Signal Amplification (TSA) (Opal Kit, Perkin Elmer, Waltham, MA, USA) according to the manufacturer’s instructions to detect. The sections were processed and the images were acquired using the Vectra 3.0 Automated Quantitative Pathology Imaging System and analysed using InForm software (v2.4.1, Perkin Elmer, Waltham, MA, USA). An average of 11 ROIs per tissue section were utilized to perform the analysis. As the analysis of the study is based on using algorithms to differentiate between the markers, we performed a thorough initial optimization to avoid false double positives. This was at first achieved by optimizing the dilution of the antibodies at different concentrations and on different channels. We found that Vista (1:500, Opal 520), CD4 (1:1000, Opal 570), PanCK (1:200, Opal 620), Iba-1 (1:50000, Opal 650), CD8 (1:1500, Opal 690) gave us the best distinction with no bleed over of the channels. We also trained the InForm software, to avoid nuclei which could be colocalizing and hence ensuring to circumvent any false double positives. Furthermore by using sections of only 4 um in thickness aided us in bypassing colocalized nuclei as usually one nucleus is about 5-6 um.

IHC was also performed on four-micron whole sections using the MACH1 Universal HRP-Polymer Detection Kit (BioCare Medical, Pacheco, CA, USA) according to the manufacturer’s instructions. Briefly, the sections were dewaxed, de-paraffinized and rehydrated in decreasing concentrations of alcohol (100-70%). Heat-induced antigen retrieval was performed using sodium citrate buffer (0.01M, pH 6.0) at 110°C for 10 mins in a decloacking chamber (BioCare Medical). The sections were treated with 0.3% Hydrogen peroxide for 30 min to remove endogenous peroxidases and then with MACH1 Sniper blocking reagent (BioCare Medical) to avoid non-specific antibody staining. Primary antibody diluted in DaVinci Green Diluent (BioCare Medical) was applied to the sections and the slides were incubated for 3.5 hrs at RT in a humidified slide chamber. MACH-1 anti-rabbit secondary antibody conjugated to horseradish peroxidase was applied for 30 min at RT followed by the Diaminobenzidine (DAB) chromogen substrate for 5 min. The slides were counterstained with Hematoxylin for 4 min and cover-slipped using the DPX mountant (Sigma Aldrich, St Louis, MO, USA). For analysis, the slides were scanned using the Aperio AT Turbo (Leica Biosysems, Wetzlar, Germany) at 40x magnification.

### Dataset analysis

Raw gene counts were initially filtered, removing any gene with less than 1 count per million in at least 2 samples. The raw data was subsequently normalized using the trimmed mean of M values (TMM) approach using the edgeR package, resulting in an expression matrix with 226 samples and 18735 genes. To rank the expression of VISTA, genes within each sample were sorted according to their expression; with the genes having the highest expression receiving the highest rank and vice versa for the gene with the lowest expression. To account for ties in rank between genes, if two or more genes had the same rank, then, their rank was replaced by the average of their initial rank. To perform the ranking, the rank function in base R was used, with default parameters.

### Statistical analysis

All statistical analysis and the preparation of graphs were achieved by using GraphPad Prism software(v8.2). The data were analyzed using non-parametric t-tests with p-values <0.05 considered to be significant.

## 2. Results

### Cohort description and image cytometry

In this selected cohort, the median age at which the primary breast cancer was diagnosed was 45 years, with brain metastasis diagnosis at 47 years; median time to develop brain metastasis was 25 months from primary diagnosis. Half of the analysed patients within this study (53%) had triple-negative BC - (TN)) followed by HER2+; 34% and ER+; 13% (Figure-1a). These breast cancers were also high grade with 56% patients falling into grade 3 category and 24% in grade 2 (Figure-1b). The tumor infiltrating lymphocytes were scored according to the International TILs working group [21], and we restratified the groups into TILs density criteria based upon (>30%, high or <30%, low) (Figure-1c) This segregation was significantly prognostic (p<0.01) for brain metastasis specific survival (BrMSS) with a hazard ratio of 2.9 (Figure-1d).

**Figure 1.**
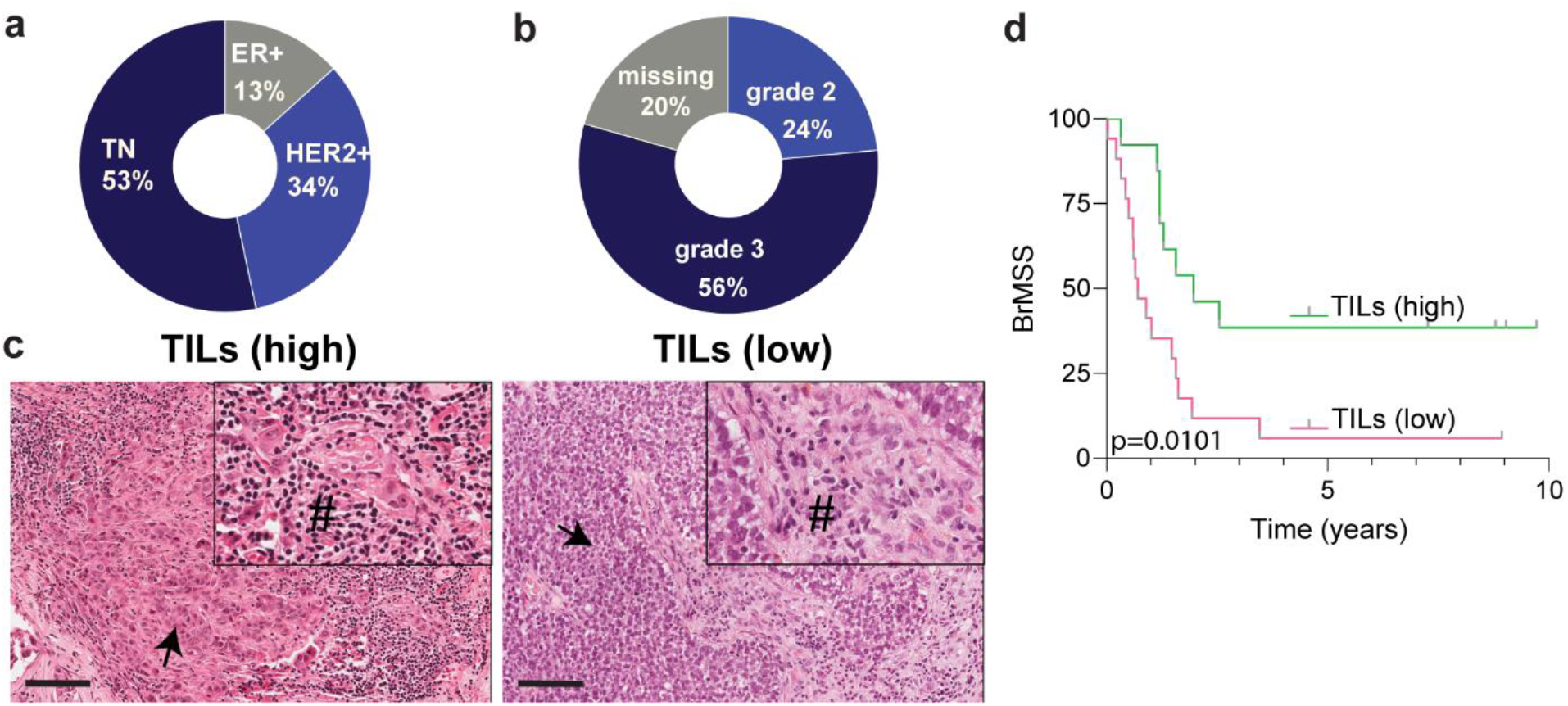
Cohort demographics and survival analyses. (**a**) Subtype breakdown of the primary breast cancers giving rise to the brain metastases; (**b**) Tumor grade breakdown of the cohort; (**c)** Hematoxylin and eosin stained representative image for TILs scoring and grouping; (**d**) Kaplan-Meier curve analysis showing brain metastasis specific survival of the TILs high and TILs low groups; # *TILs; arrows indicate tumor. Scale bar 100 μm*.

We performed transient knockdown of *VSIR* gene on the breast cancer cell line MDA-MB-468 to test the specificity of VISTA antibody. Antibodies against the other markers were extensively used and well characterised hence, eliminating the need to re-validate them. As demonstrated in Figure 2a, at high and low magnification we observe positive staining for VISTA protein in the cells that received scrambled control sequence. Contrastingly, in the cells that received the short interfering RNA sequence against *VSIR* we observe no staining at the protein level. After validation of the antibody, we performed multiplex immunofluorescence (mIF) labelling using tyramide signal amplification (TSA) to profile CD4, CD8, VISTA, Pan cytokeratin (PanCK) and Iba-1 positive cell subsets within this cohort. Images were acquired on the Vectra automated quantitative digital pathology platform and data was processed using InForm software’s standardized workflow tool. Spectral unmixing precisely separated the staining patterns of each different protein by employing the distinctive features of the dyes and was performed to ensure no bleed through or overlapping of the adjacent fluorochromes (Figure-2b) thus limiting false positive results. Representative images of the separation of DAPI, PanCK at 620 nm, CD4 at 570 nm, Iba-1 at 650 nm, CD8 at 690 nm, and VISTA at 520 nm are shown in Figure-2b; a clear distinction of the cellular subsets is observed at the different wavelength channels for each marker. Following tissue and cell segmentation, we integrated the respective segmentation dataset employing the FCS express 6 cytometry software. This enabled us to analyze the multiplexed images from 10-15 randomly selected regions of interest for each case (Figure-2c). We quantified the multi-parameter spatial (different regions of the tumor)-as well as single-cell cluster phenotypes within our dataset. The data was at first scrutinized at the tissue level with tumor and tumor-associated area/stroma separated on a histogram (Figure-2c). We also visualized the data on histograms for each tissue region to separate the CK+ (tumor/epithelial cells) and CK-(non-tumor cells), followed by CD8+ and CD4+ subsets (Figure-2c). Approximately 15 000 cells per case were analyzed per case (Figure-2d).

**Figure 2.**
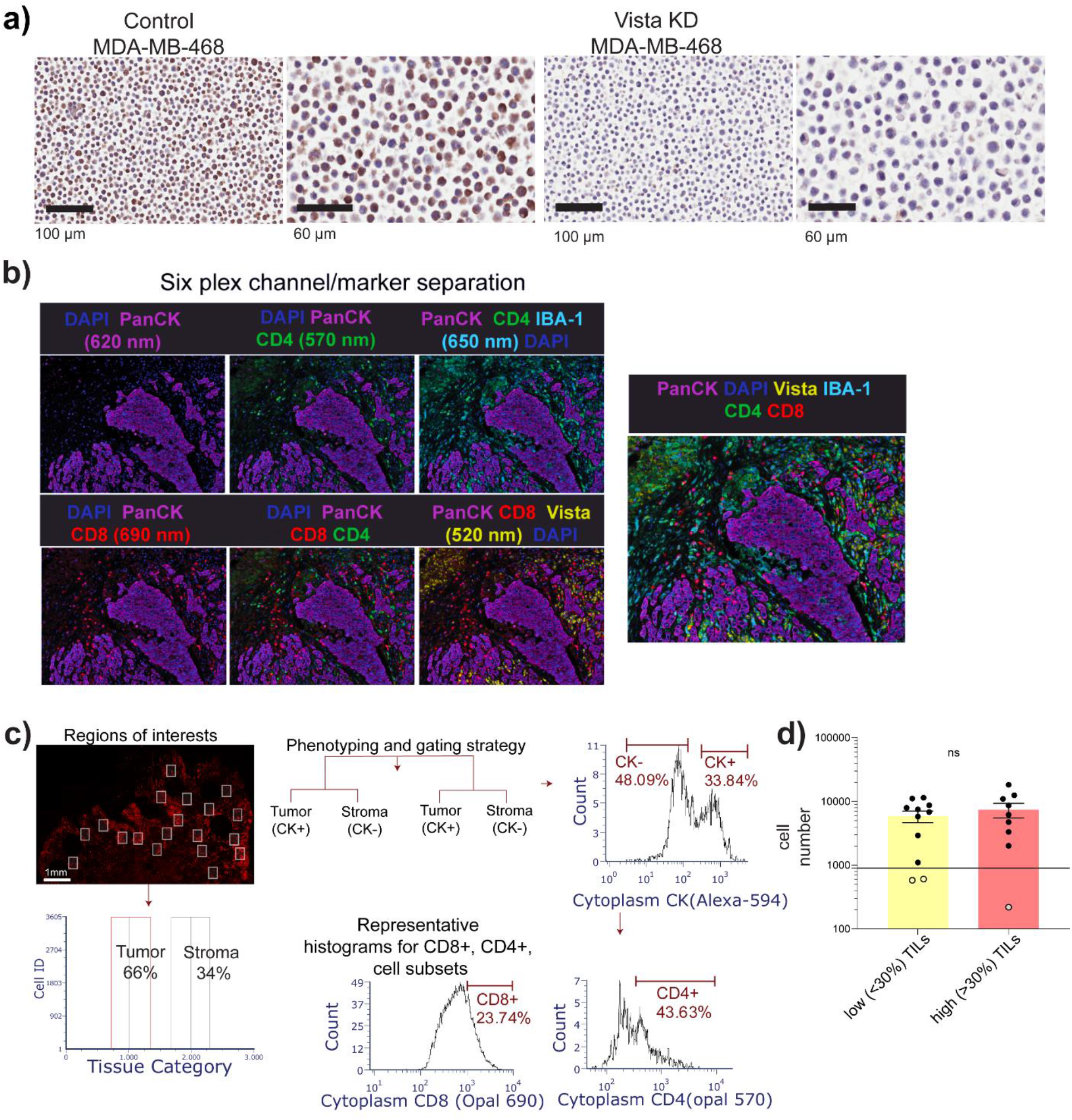
Imaging, tissue, and cell segmentation. **a**) VISTA antibody validation on MDA MB 468 cells using IHC using scrambled control and siRNA against *VSIR* gene b) Spectral unmixing and separation of the fluorochromes highlighting the different cell subsets; (**c**) Data integration and representative gating strategy for single-cell subset analysis in different regions of the samples; example of the ROIs in a case followed by separation of the tumor and stroma regions based on PanCK expression. These parent cell populations were further gated for CD8, CD4+ cell subsets (d) Average cell numbers from each group being analysed in the study; each dot is representing a case.

### Increased VISTA and Iba-1 expression in the TME

We segmented the tissue regions into tumor and stroma and analysed each compartment to delineate cellular phenotypes. We considered cells showing positive expression of PanCK to be tumor epithelial cells and quantified the subset of TILs as indicated in Figure 2c. Initially we considered single marker positive cell subsets and found that VISTA was expressed on the tumor cell surface in all samples however, the proportion of VISTA+ tumor cells differed between the groups (Figure-3a).The percentage of tumor cells that expressed VISTA on their surface was 50% in the low TILs group compared to 23% in the high TILs group (p=0.0027; Figure-3a).

**Figure 3.**
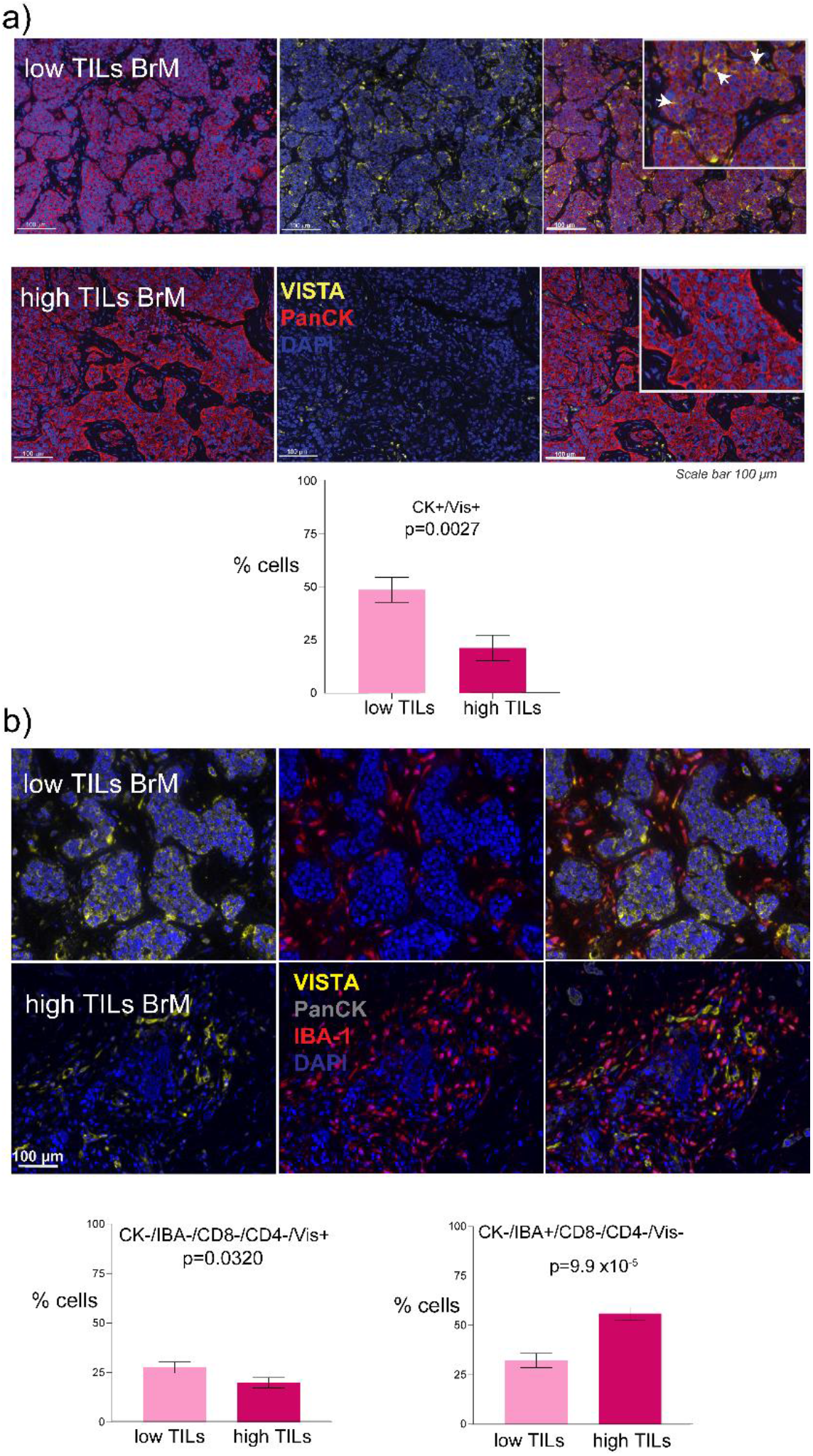
VISTA and Iba1+ cellular subtypes a) VISTA expression on the tumor cell surface of the low *vs.* high TILs groups and the associated bar graph; arrows indicate vista positive tumor cells (b) Images represent the expression of VISTA and Iba1+ cells and bar graphs showsVISTA and IBa1+ exclusive cells Between the low and high TILs groups respectively.

We also found its exclusive expression in 27% of the low TILs group, which was significantly higher than 19% in the high TILs group (p=0.0320; Figure-3b). Interestingly, VISTA has been reported to be highly expressed by microglia and especially differentially expressed in central nervous system (CNS) pathologies [22]. Therefore, we next asked if there was an association between VISTA and the microglial marker Iba-1 expression in the TME. VISTA and Iba-1 co-expression was found to be present in about 25% of the microglia (Suppl Figure-1i). Interestingly, 40% of both groups displayed negativity for all the markers used in the study (Suppl Figure-1ii). As the presence of microglia in the brain TME has been previously described [23] and we also investigated if there was a difference in the proportion of microglial cells between low and high TILs groups. We observed that 60% of the TME cells were Iba1+ in the high TILs group (p=9.9 x 10^-6^; Figure-3b) compared to 30% in the low TILs group.

### Phenotypes of the CD8+ T cells

We investigated the CD8+ and CD4+ T-cell phenotypes in our cohort. The CD4+ T cells did not differ significantly in their proportion in low and high TILs groups, 30% of CK-cells were CD4+ in the low TILs and 38% in the high TILs group (Suppl Figure-1iii). We also found 37% and 40% of CD4+ cells co-expressed Iba-1 (Suppl Figure-1iv). Double positive subsets within CD4+ T cell population such as Iba-1+/VISTA+ or IBA-1+/VISTA-were found to be similar in their proportions in both the groups (Suppl Figure1v, vi, vii, viii). In tumors with high and low TILs (Figure-4a), 25% and 26% of CK-cells were CD8+ T-cells respectively (Figure-4b). Interestingly, this changes for double positive cells; where 46% of CD8+/Iba-1+ are found to be present in the high TILs groups compared to 26% in the low TILs (p=0.0110; Figure-4c). We investigated if this was true when triple positive populations were analysed and this pattern remained the same with high TILs group displaying higher (49%) CD8+/Iba-1+/VISTA-population (p=0.0463; Figure-4Aiv). When we examined VISTA expression within CD8+ population (Figure-4e) we found 30% of CD8+ T cells express VISTA in the low TILs group compared to 18% in the high TILs group (p=0.0080; Figure-4f). Intriguingly, when we investigated Iba-1 positivity within CD8+/VISTA+ population, we found a similar trend with CD8+/Vis+/Iba-1+ or Iba-1-cells; where low TILs group had higher amount of both populations 44% (p=0.0021; Figure-4g) and 26% (p=0.0092; Figure-4h) compared to high TILs tumors respectively.

**Figure 4.**
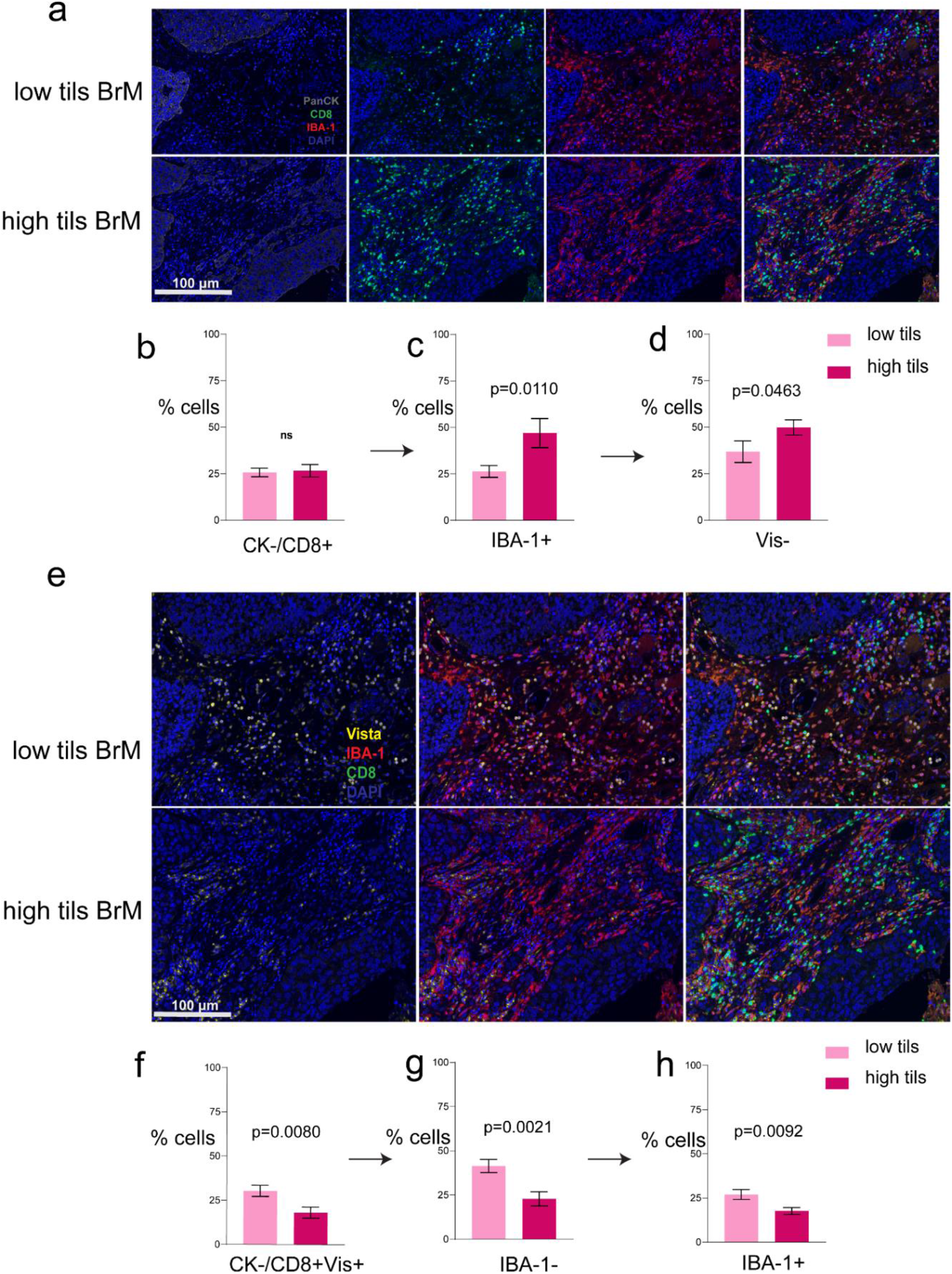
CD8+ T cell populations with arrows indicating the direction of the subsequent gated cell populations within the parent population a) Representative images of mIF:CK, CD8, and Iba-1; b) CD8+ gated population of cells comparison between the low and high TILs groups; c) Bar graphs showing Iba-1+ cells within the CK-/CD8+ parent population; d)VISTA negative cells within the CK-/CD8+/Iba-1+ cells; e) Representative images of mIF: CD8, VISTA and Iba-1; f) CK-/CD8+ gated parent population with VISTA expression; g) Bar graphs showing Iba-1-cells within the CK-/CD8+Vis+ parent population; h)Iba-1+ cells within the same parent population

### Validation of mIF using IHC and RNA expression analysis

After quantifying the phenotype of the TILs using mIF, we next attempted to validate our findings using standard immunohistochemistry of VISTA, Iba-1, CD4 and CD8 on eight randomly selected whole sections; four from low TILs and four from high Tils within our cohort. Upon semi-quantitative scoring, Vista was positively stained on the tumor cells for three out of four cases in the low TILs group. In the high TILs tumor no tumor cell positivity was evident on single IHC for VISTA (Figure-5a). Iba-1 (Figure-5a) was also found to be highly expressed in the TME of high TILs BrMs compared to low TILs (Figure-5a). We also observed similar pattern of positive staining for CD8 and CD4 in the TME in low and TILs tumors with no evident difference (Figure-5a). To further delineate the expression of VISTA in BrMs, we explored the Klemm dataset [12], where after flow cytometry assisted separation of immune cell and non-immune cells, they performed RNA-sequencing on BrMs and glioma samples. We examined the expression of *VSIR* in BrMs and gliomas in the non-immune (CD45-) population and found BrM non-immune cells to have a significantly higher level of VISTA expression (p=0.0453; Figure-5b). Ranking of VSIR gene within the CD8+ population in BrMs revealed that in 75% of cases, VISTA’s abundance is in the top 10 percentile (top quartile) of all ~18000 genes expressed in this cell subtype (Figure-5c). We also compared the expression of *VSIR* in the microglial (MG), monocyte-derived macrophages (MDMs) and non-immune population (CD45-) within BrMs. Interestingly, we found that in BrMs *VSIR* expression was significantly higher in the cells from myeloid lineage, especially microglia (Figure-5d). This population had significantly higher expression of *VSIR* compared to CD8+ (p=0.043; Figure-5d) as well as CD45-cells (p<0.000001; Figure-5d). The expression levels were comparable between the MDMs and CD8+ cells, however, *VSIR* expression was higher in MDMs compared to CD45-cell (p<0.000001; Figure-5d).

**Figure 5.**
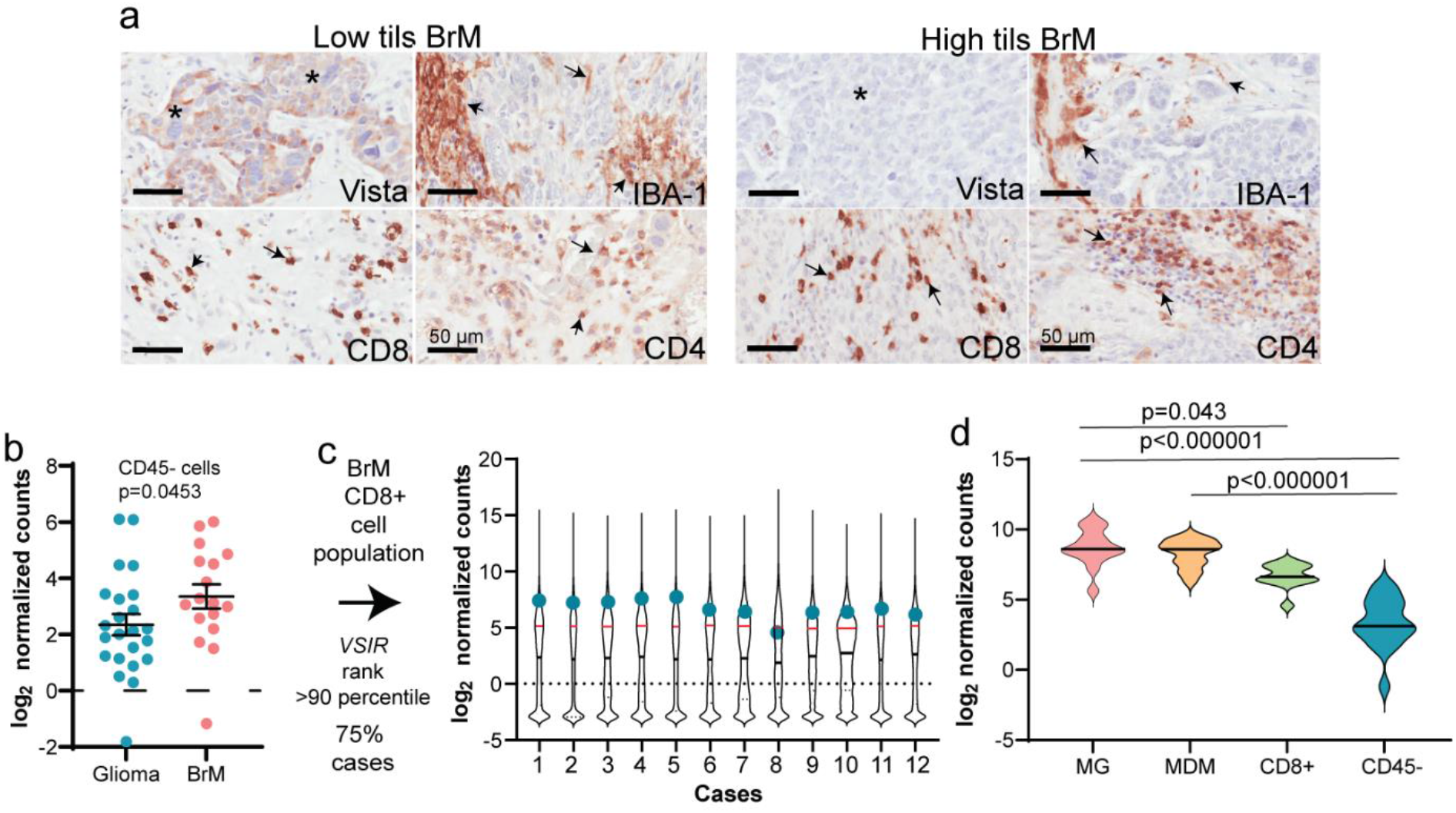
VISTA expression analysis using single IHC as well as RNA analysis in the Klemm dataset. (a) Representative images of standard IHC staining on BrMs sections with * representing tumor cells and arrows indicating positive staining. (b) *VSIR* gene expression in BrMs and Gliomas within CD45-population in the Klemm RNAseq dataset [12]; (c) Ranking of VSIR in CD8+ population within BrM cases;(d) Within BrMs expression levels of VSIR in Microglial (MG), monocyte-derived macrophages (MDMs), CD8+ and CD45-populations.

## 3. Discussion

The brain TME is a rapidly evolving field of research and its importance is widely recognized as a critical regulator of progression and treatment resistance. In spite of emerging data associating tumor infiltrating lymphocytes to prognosis in BrMs [24–26], the immune cell subsets and phenotypes within BrM are not well-defined. We analyzed by employing a mIF approach, a targeted panel of immune-related proteins; CD4, CD8, Iba-1, and VISTA which have not yet been described in brain metastasis in either the single- or co-expression context. We specifically investigated the CD8+ and CD4+ T cells and compared their phenotypes in BrMs with high and low infiltrating lymphocytes which were also prognostic. We interrogated how the TILs in patients with better prognosis differ in their phenotype compared to the ones with poor prognosis within our selected cohort.

Our data showed that VISTA is highly expressed on the tumor cell surface of BrMs and this was significantly higher in the group with low density of TILs. Overall, we observed an enhanced expression of VISTA in the tumor compartment as well as in the microenvironment of patients with low density of TILs. Interestingly these patients also had a poorer prognosis compared to the high TILs group. In this setting, VISTA might be manifesting immune-modulatory effects, especially in tumours with low density of TILs. However, we cannot eliminate the possibility that VISTA expression on the tumor cell surface may have caused less TILs to infiltrate the area.

VISTA is a negative immune checkpoint regulator, is highly expressed in myeloid cells and its expression is known to be suppressive on resting as well as activated CD4+ and CD8+ T-cells [27]. Our findings on VISTA being expressed by epithelial cells confirms data in other cancers [19,27–31]. In non-small cell lung carcinomas (NSCLC), VISTA was found to be expressed in majority of NSCLCs and contrastingly this expression associated with lymphocyte infiltration and PD-L1 expression [28]. In ovarian cancer, expression of VISTA has contradicting reports. On one hand two studies indicate VISTA to be highly enriched in tumors leading to a poor prognosis and advanced disease and on the other, one study has been shown VISTA to be associated with good prognosis. Mulati and colleagues found VISTA to be expressed in 91% of the samples and in an animal model bearing ovarian tumours, anti-VISTA antibody increased the longevity of the treated animals by reducing the tumor burden. In-vitro studies showed that tumor cell expression of VISTA caused immune evasion by suppressing T-cell proliferation and cytokine production [27]. Additionally, VISTA expression has been linked to the progression of the disease in ovarian cancers. The authors noted that expression of VISTA increased with advanced disease and lymph node metastasis [19]. Contrastingly, VISTA’s expression has also been shown to associate with a favorable prognosis in a different cohort of high-grade ovarian cancers. This retrospective study had used tissue microarrays to assess the expression of VISTA on tumor cells as well as in immune cells [31]. These two studies had different scoring criteria, and the study by Liao and colleagues used whole sections which might have provided a higher accountability for the heterogeneity within the section. Nevertheless, more in-depth analysis is required to understand the contrasting associations of VISTA in ovarian cancer. Our observation of co-expression of VISTA on CD8+ T cells on low TILs density can also be extrapolated to suggest that perhaps VISTA expression is causing immune evasion.

With respect to Central Nervous System metastatic disease, recent work by Guldner and coworkers have identified brain resident myeloid cells to be inducing an immunosuppressive microenvironment, thus promoting metastasis [20]. This study generated mouse models of BrMs to derive functional understanding of the myeloid cell population in this disease. When they performed mechanistic in-vitro and in-vivo studies they found C-X-C motif chemokine ligand 10 (Cxcl10) promoted BrMs, and as this chemokine is an immune modulator, they further characterized to elucidate the mechanism behind it. Investigation of publicly available human CNS myeloid RNAseq datasets, they found increased expression of Vsir (VISTA) and CD274 (PD-L1) in the myeloid cells [20]. In addition, they found co-inhibition of PD-L1 and VISTA in a mouse model of brain metastasis reduced the metastatic burden. Our results are complimentary to their finding where we have shown increased VISTA in a sub-population of patients with BrMs and its expression is an indicator of poor prognosis. Furthermore, we observe an enhanced expression of VISTA in BrMs compared to gliomas as well as VISTA was more pronounced in the myeloid lineage cells in our mIF study as well as in the Klemm dataset [12]. To our knowledge our study is the first to show this differential expression of VISTA in the tumor as well as in CD8+ T cells in a BrM clinical cohort.

Brain resident macrophages/microglia perform numerous roles in brain health and disease [32]. With respect to BrMs, microglia are known to be multifaceted in their function such as promoting proliferation, angiogenesis and invasion in BrMs [33]. Recent work by Simon and co-workers, highlighted using intravital microscopy how microglia are recruited and activated after xenotransplantation of breast cancer cells into the brain [34]. This study showed that microglial activation and accumulation around the lesion can cause infiltration of immune cells as well as aberrant electrical activity in the brain. Similarly, in a mouse model of BrM, microglial accumulation has been associated with increased tumor burden. When selective depletion of the anti-inflammatory microglial phenotype was carried out, a reduction in tumor burden was observed [35]. These studies highlight the importance or microglia in brain metastasis. For this study we used a common microglial marker Iba-1 in our panel to investigate the expression levels and associations of Iba-1+ microglia.

Our findings suggest increased microglia in the tumours with high density of TILs and we also observe increased CD8+ T-cells that co-express Iba-1. Simultaneous accumulation of microglia and infiltration of immune cells has been shown in a mouse model of BrM [34]. Although no mechanism is known for this association, this accumulation was found to reduce the density of tumor cells in the brain [34]. We observed increased microglia in the high TILs group which also had a better prognosis. This suggests that perhaps increased microglia may have caused anti-tumour immunity. Additionally, our perplexing finding of double positive CD8+/Iba-1+ cells, leads us to suggest that these double positive cells may be representing pro-inflammatory M1 macrophages. This is plausible because Boddaert and colleagues used a rat model for stroke to demonstrate increased CD8 expression in activated macrophages using Iba-1 as a marker. Their results corroborate our finding of double-positive macrophages. Furthermore, in-vitro analysis revealed CD8 stimulation caused repolarization of M2 to M1 phenotype along with highly proliferative Iba-1+CD8+ cells [36]. If this is true for our findings, then this co-expression could play a protective role in BrMs. It is known that microglia can exist in two states, anti-inflammatory which facilitates invasion, angiogenesis and tumour growth [33,37]; and pro-inflammatory which can release proinflammatory cytokines, eliciting a T-cell response [38]. Pro-inflammatory microglial phenotype switch can infer a microglial population which may have caused anti-tumor immune response[39]. Nonetheless, we cannot rule out the possibility of false double positive cells, however we strongly believe this is highly unlikely. Future studies are required to investigate a larger population of resected tumors using flow cytometry to see if double positive CD8+/Iba-1+ are a consistent phenotype in BrM.

We appreciate the limitations of this study such as the size of the cohort which needs to be expanded to further validate our findings. We have used commercially validated antibodies however, batch variability might still play a role and therefore, future studies should focus on employing various commercially available antibodies to test the suggested phenotypes of TILs in BrM from this body of work. In addition, we cannot eliminate the effect of treatment that may be impacting the TILs phenotype as our patient population ranges from early to late 2000s and the treatment regimen has drastically changed within the last decade. For example, it would be crucial to test Vista expression in a patient population treated with current immunotherapy regimens as that might provide novel insights into treatment response. Furthermore, BrMs arising from different primaries may have different phenotypes which need to be further elucidated. Finally the functional implications of these findings using different in-vitro and in-vivo models would be essential in the future for clinical translation.

Overall, we have utilized mIF as a tool to provide a proof of concept study to demonstrate TME distinctive changes in the TILs of BrM. Through this we have defined the expression of a new immune checkpoint molecule in a clinical cohort of BrMs.

## 5. Conclusions

Brain metastasis from the breast is one of the common causes of mortality in women, the poor quality of life and the associated morbidity makes it compelling to understand the disease better. An emerging and evolving concept within the brain TME is the existence of the complex landscape of cells within this unique organ. The positive brain tumor responses to immune checkpoint inhibition raises optimism that extensive understanding of the biology of the tumor will lead to better management. There is hope that developing a comprehensive understanding of the TME will lead to design of more effective immunotherapies. Our study has, shown a novel immune checkpoint, VISTA, to be highly expressed in brain metastasis and its expression is higher in the group of patients who have a worse prognosis compared to the high TILs group. We have also provided insights into the VISTA co-expressing immune cell subsets within our selected cohort. We found increased microglia to be more pronounced in BrMs with high density of TILs and we also observed CD8 + Iba-1 cells. Further studies are required to elucidate the mechanisms of differential expression of VISTA and microglia in BrMs.

## Supplementary Materials

**Figure S1.**
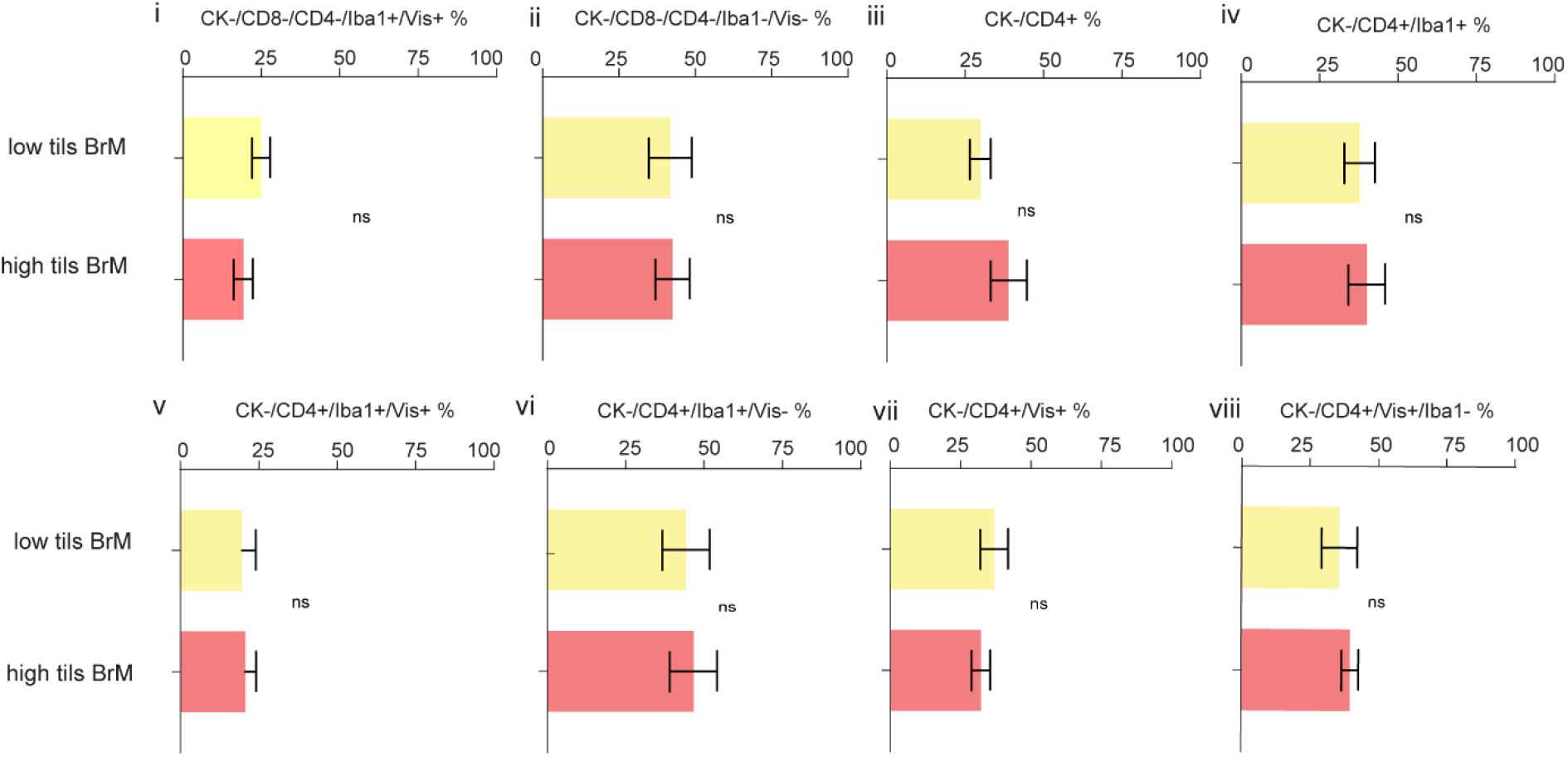
Quantified bar graphs of CD4+ subsets within the groups.

## Author Contributions

Conceptualization, PkdC and PS; methodology, PKdC, HC; software, PKdC, TN,; validation, PKdC, HC, ML; data curation; PKdC, ML, XDL, KF, CN, WGK, JMS; writing—original draft preparation, PKdC; writing—review and editing, All authors; supervision, SRL.; project administration PKdC and SRL. All authors have read and agreed to the published version of the manuscript.

## Acknowledgments

The authors thank Mr Clay Winterford for his histotechnological expertise. We are grateful for support from Metro North Hospital and Health Service and we thank the patients past and present who donate tissue and clinical information for research.

